# Control of iron homeostasis by a regulatory protein-protein interaction in *Bacillus subtilis*: The FurA (YlaN) acts as an antirepressor to the ferric uptake regulator Fur

**DOI:** 10.1101/2023.09.28.559918

**Authors:** Lorenz Demann, Rica Bremenkamp, Kolja Stahl, Björn Hormes, Robert Warneke, Juri Rappsilber, Jörg Stülke

## Abstract

Iron is essential for most organisms. However, two problems are associated with the use of iron for aerobically growing organisms: (i) its accumulation leads to the formation of toxic reactive oxygen species and (ii) it is present mainly as the highly insoluble ferric iron which makes the access to iron difficult. As a consequence, a tight regulation of iron homeostasis is required. This regulation is achieved in many bacteria by the ferric uptake repressor Fur. The way how the activity of Fur is controlled, has so far remained elusive. Here, we have identified the Fur antirepressor FurA (previously YlaN) in the model bacterium *Bacillus subtilis* and describe its function to release Fur from the DNA under conditions of iron limitation. The FurA protein physically interacts with Fur, and this interaction prevents Fur from binding to its target sites due to a complete re-orientation of the protein. Both *in vivo* and *in vitro* experiments using a reporter fusion and Fur-DNA binding assays, respectively, demonstrate that the Fur-FurA interaction prevents Fur from binding DNA and thus from repressing the genes required for iron uptake. Accordingly, the lack of FurA results in the inability of the cell to express the genes for iron uptake under iron-limiting conditions. This explains why the *furA* gene was identified as being essential under standard growth conditions in *B. subtilis*. Phylogenetic analysis suggests that the control of Fur activity by the antirepressor FurA is confined to, but very widespread in bacteria of the class Bacilli.

**IMPORTANCE:** Iron is essential for most bacteria since it is required for many redox reactions. Under aerobic conditions, iron is both essential and toxic due to radical formation. Thus, iron homeostasis must be faithfully controlled. The transcription factor Fur is responsible for this regulation in many bacteria; however, the control of Fur activity has remained open. Here we describe the FurA protein, a so far unknown protein which acts as an antirepressor to Fur in *Bacillus subtilis*. This mechanism seems to be widespread in *B. subtilis* and several important pathogens and might be a promising target for drug development.

## INTRODUCTION

Living organisms require the presence of micronutrients to survive. Iron is one of these micronutrients that is essential; however, trace amounts of the metal are sufficient to sustain the life of bacteria. The difference between iron and other trace minerals such as zinc or manganese is the facile redox transformation between ferric (Fe^3+^) and ferrous iron (Fe^2+^) (1). Enzymes that bind iron as a cofactor make use of this transformation for electron transport with the iron acting as the electron acceptor and donor.

As many other essential metal ions, iron is not only required for life, but its accumulation can also have toxic effects. During aerobic respiration superoxide radicals are produced as byproducts. To remove superoxide, it is converted to H_2_O_2_ by the enzyme superoxide dismutase. H_2_O_2_ can then be detoxified by catalase or peroxidase but it can also react with ferrous iron via the Fenton reaction. A product of this reaction is the hydroxyl radical (for review see 2, 3). H_2_O_2_ can also react with superoxide in the ferrous iron catalyzed Haber-Weiss reaction to yield the hydroxyl radical (4). These radicals can cause damage to many biological macromolecules which can ultimately lead to cell death (5). To prevent such damage, cells can detoxify superoxide and H_2_O_2_, repair organic molecules such as DNA, and finally regulate the amount of free ferrous iron in the cell.

Life on earth emerged under anaerobic conditions; thus iron was mostly present in the highly soluble ferrous form. Presumably, bacteria adapted very early to the use of iron for enzymatic reactions (6). With the development of oxygenic photosynthesis, the atmosphere turned aerobic, and this drastically changed bacterial evolution with respect to iron metabolism. On one hand, bacteria evolved aerobic respiration because the oxidation power of oxygen ultimately yielded much more energy than fermentation or any other respiration. On the other hand, this also led to the probably unintended production of superoxide and H_2_O_2_. As described, both compounds readily react with ferrous iron to form reactive oxygen species. Simultaneously, with increasing amounts of oxygen in the atmosphere ferrous iron rapidly oxidizes to the highly insoluble ferric iron, or more specifically to ferric hydroxides that rapidly precipitate (from 10^-6^ M free ferrous iron under anoxic conditions to 10^-10^ M complexed and 10^-17^ M free ferric iron under aerobic conditions (7). Bacteria require 10^-5^ to 10^-7^ M for optimal growth, so they had to develop mechanisms to acquire sufficient iron from their environment (1).

Iron acquisition is growth-limiting for many bacteria, in particular for pathogens that acquire iron from the host. To make insoluble ferric iron available for the cell, bacteria use siderophores, low molecular weight compounds that bind ferric iron with high affinity and thereby solubilize it (1). The ferri-siderophore complex is then taken up by specific transport systems, typically ABC-transporters in Gram-positive bacteria (8). The Gram-postive model organism *Bacillus subtilis* encodes proteins for the synthesis of several siderophores and systems for the uptake of different iron sources (9). Additionally, many bacteria also produce an uptake system for elemental iron (10, 11). In *B. subtilis*, this transporter is named EfeUOB and transports elemental ferric iron into the cell (12).

Since iron is essential but also toxic, intracellular iron levels need to be tightly regulated. This regulation is mediated by Fur, the ferric uptake regulator. According to the COG database (13), Fur is conserved in many archaea and most bacteria. Fur belongs to the ferric uptake regulator family which includes Zur, the major regulator of zinc homeostasis and PerR which responds to peroxide stress. Fur regulates the expression of at about 60 genes in *B. subtilis* (14, 15, see http://www.subtiwiki.uni-goettingen.de/v4/regulon?id=protein:F899F1EE27E6D503BCC06BC52E3C7FD80B8EF725 for a complete list of the *B. subtilis* Fur regulon, 16). Fur almost exclusively represses the expression of its target genes which are mostly iron uptake and siderophore synthesis systems (14). An exception is the Fur-activated *pfeT* gene which encodes an efflux pump (17, 18). Fur consists of an N-terminal DNA-binding domain and a C-terminal domain which is thought to bind ferrous iron as a cofactor and which mediates dimerization (19, 20). It has long been assumed that Fur binds ferrous iron as a cofactor but only recently it could be shown that Fur actually reversibly binds an iron sulfur cluster in *Escherichia coli* (21).

Even though it could not be shown, it is widely assumed that *B. subtilis* Fur senses the intracellular iron concentration by binding ferrous iron (22, 23). With decreasing extracellular iron concentrations the Fur regulon is derepressed in three waves in *B. subtilis* (15). First, iron uptake systems for elemental iron, ferric citrate and petrobactin are expressed. Sequentially, the synthesis of the siderophore bacillibactin and uptake systems for bacillibactin and the hydroxamate siderophores to scavenge iron occurs before the iron-sparing response is initiated to inhibit the translation of iron binding proteins mediated by the regulatory RNA *fsrA* (24).

Despite the absence of direct evidence for the binding of iron to Fur, there has been only little research regarding possible other mechanisms that regulate Fur (25). However, recent studies revealed that there are proteins which modulate the activity of Fur in different bacteria. In *Salmonella enterica*, the EIIA^Ntr^ protein of the non-canonical phosphotransferase system regulates the expression of iron uptake genes via Fur by a direct protein-protein interaction which results in the release of Fur from its DNA binding sites (26). Another but similar mechanism was recently found in uropathogenic *E. coli*, where the proteins YdiV and SlyD cooperatively bind Fur and reduce its DNA binding (27). These results indicate that the regulation of Fur, which was assumed to be exclusively dependend on ferrous iron, actually involves Fur antagonists that might be more common then previously anticipated.

We are interested in the functional characterization of unknown or poorly studied proteins in the model bacterium *B. subtilis*. Although *B. subtilis* is one of the best studied bacteria, the function of many proteins remains to be elucidated. The YlaN protein a highly abundant but only poorly studied protein (28, 29). Moreover, the *ylaN* gene is essential under standard growth conditions (30, 31). Both the very high expression and the essentiality suggest that the YlaN protein plays a very important role in the cell. Recently it was shown that t he *ylaN* gene becomes dispensable when ferric iron is added to the growth medium (32). This provides a strong indication that the YlaN protein is involved in the control of iron homeostasis. Moreover, recent *in vivo* crosslinking data revealed an intriguing interaction between YlaN and Fur which indicates that YlaN might be another Fur antagonist and thus exert its role in iron homeostasis via Fur (33, 34).

In this work, we have studied the role of YlaN in the control of iron homeostasis in *B. subtilis*. We confirm the direct protein-protein interaction between Fur and YlaN, and show that Fur is unable to bind its target DNA in the presence of YlaN. Thus, YlaN acts as an molecular antirepressor of Fur which we rename FurA.

## RESULTS

### FurA becomes dispensable in the absence of Fur

It has been shown that the essential *furA* (previously *ylaN*) gene becomes dispensable if ferric iron is added to the growth medium (32). Based on the interaction between FurA and the Fur regulator protein, we hypothesized that FurA might antagonize Fur to allow the expression of iron uptake systems at low iron concentrations. If this were true, the deletion of the *furA* gene under standard conditions might be toxic as a result of continued repression of the genes for iron uptake by Fur. To test this idea, we attempted the deletion of *furA* in the *fur* mutant GP879. As a control, we used the isogenic wild type strain *B. subtilis* 168. In agreement with the published data, *furA* could not be deleted under standard conditions or if the plates were supplemented with ferrous iron. However, the deletion was possible if the medium was supplemented with ferric iron. In contrast, the *furA* gene could be deleted under all three conditions in the *fur* mutant strain GP879. These results confirm the conditional essentiality of FurA depending on the iron supply and they suggest that FurA might be needed to prevent some harmful activity of Fur under conditions of iron limitation.

### FurA is quasi-essential under standard growth conditions

When the concept of essentiality was introduced, a gene was regarded essential if it could not be activated under standard growth conditions for the organism, *i. e*. on LB medium supplemented with glucose at 37°C for *B. subtilis* (31, 35). Today, we know that several essential genes can in fact be deleted under standard conditions but that the mutant strains immediately acquire suppressor mutations that help to overcome the growth-limiting problem. Such genes are called quasi-essential as they remain essential under standard conditions in the genomic context of the standard wild type strain.

The possibility to delete *furA* from a *fur* mutant suggested that *furA* might also be quasi-essential. Thus, we made use of the *furA* mutant GP3324 that had been isolated in the presence of ferric iron as described above. The strain was cultivated on standard sporulation medium under iron-limiting condition. As expected, we observed the development of few individual colonies that most likely resulted from the acquisition of suppressor mutations. We re-isolated eight independent colonies that had appeared in the absence of added iron or in the presence of ferrous iron each, and sequenced their *fur* alleles as we already knew that *fur* mutants tolerate the inactivation of *furA*. Of the total of 16 mutants, all but one had single point mutations in the Fur coding sequence that resulted in amino acid substitutions in the Fur protein. These mutations all occurred in the N-terminal part of the protein which is required for DNA binding indicating that the mutated Fur versions are impaired in DNA binding (19). Several of the substitutions were observed in multiple mutants, both from seleection without added iron or in the presence of ferrous iron (see Fig. 1A). Interestingly, several of these suppressor mutants had a reddish colony colour which could also be observed in the *fur* deletion strain when grown on plates with additional ferric iron (Fig. 1B). It can thus be assumed that DNA binding of Fur in the suppressors with the reddish phenotype was completely abolished whereas DNA binding of Fur in the suppressors with white colony colour was only reduced.

**Fig. 1.**
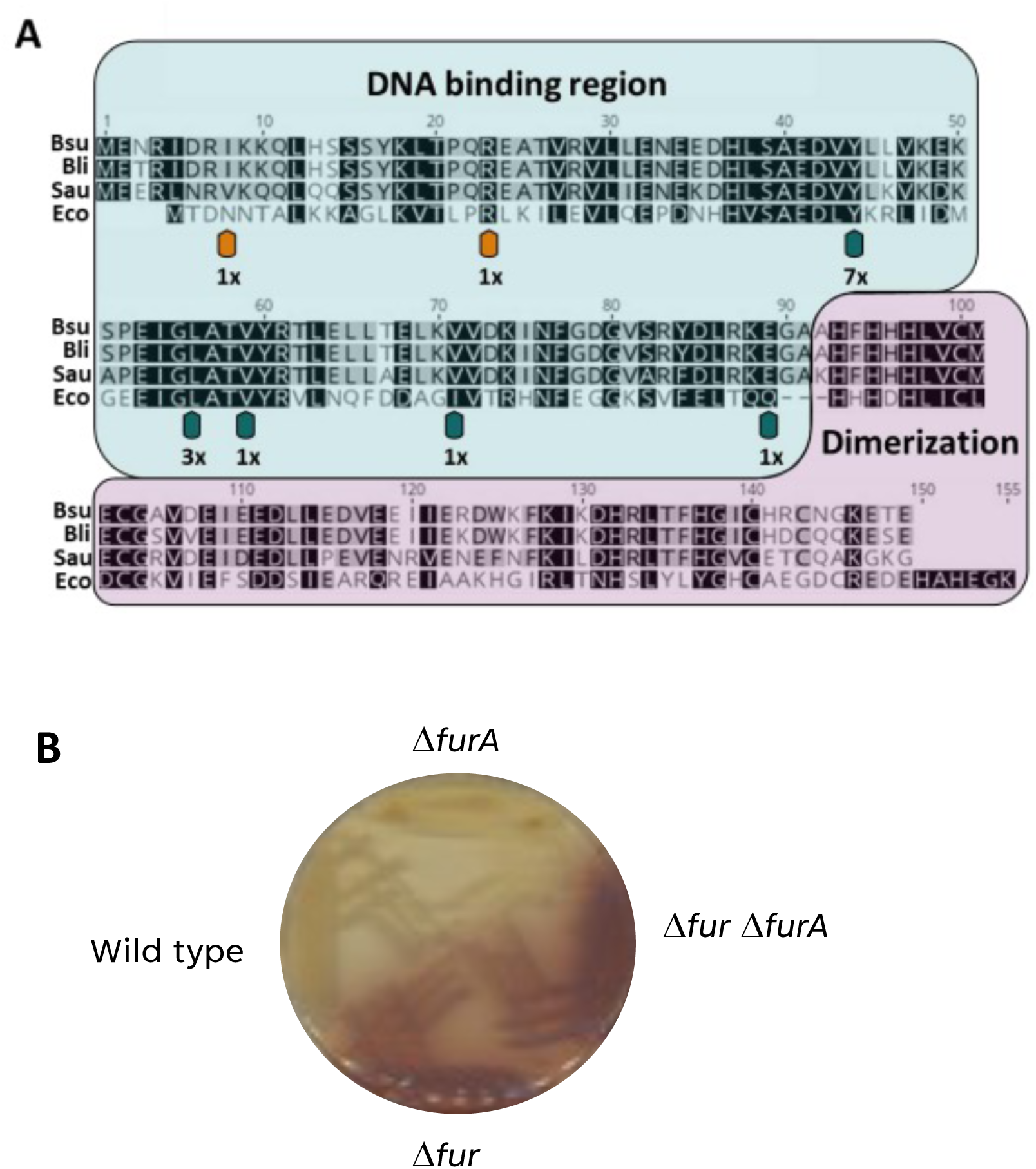
Mutations in Fur upon deletion of the *furA* gene. A. Alignment of the sequences of Fur proteins from different bacteria. The DNA-binding and dimerization domains are highlighted in light green and magenta, respectively. The positions of point mutations in the individual suppressor mutants are shown by arrows. The numbers indicate how often an individual amino acid substitution was found. The orange arrows indicate that the corresponding mutant colonies were reddish as the Δ*fur* mutant. Bsu, *B. subtilis*; Bli, *Bacillus licheniformis*; Sau, *S. aureus*; Eco, *E. coli*. B. Growth of wild type and mutant strains of *B. subtilis* on a LB agar plate containing 2.5 mM ferric iron. The deletion of the *fur* gene results in a red colony color.

One of the suppressor mutants, GP3368, had no mutation in the *fur* coding sequence. Whole genome sequencing, however, identified the accumulation of mutations in the *spoIIM-fur* region directly upstream of *fur.* Importantly, one of the mutations affects the presumptive -10 region of the fur promoter and a one base deletion results in a shortened spacer (16 bp instead of 17 bp) between the -35 and -10 regions (see Supplemental Figure S1) suggesting that the promoter was not active and that Fur was not properly expressed in this mutant.

Taken together, these results demonstrate that *furA* is a novel quasi-essential gene. Under standard conditions on complex medium, the deletion of *furA* causes the immediate acquisition of suppressor mutations that interfere with Fur activity or expression. Moreover, these results support the idea that the Fur repressor becomes toxic for the cells in the absence of FurA and iron, i.e. under conditions that require expression of the iron uptake genes that are all members of the Fur regulon.

### Ferrous iron is perceived as iron limitaton in *B. subtilis*

The attempts to delete the *furA* gene have revealed an important difference between the two different states of iron: the availability of ferric iron allowed the deletion of *furA* whereas this was not possible in the presence of ferrous iron. Moreover, cultivation of the *furA* mutant GP3324 in the absence of iron and in the presence of ferrous iron selected precisely the same mutations in Fur. Thus, the *B. subtilis* cells seemed to perceive the presence of ferrous iron as an iron limitation. To test this idea, we tested growth of the Δ*furA* strain GP3324 on plates with different iron sources. The strain was propagated directly from a cryo culture on sporulation plates with 0.5 mM of different iron sources. This concentration was used since the Δ*furA* strain can grow on plates supplemented with 0.5 mM Fe(III)citrate. No growth was observed with the ferrous iron source Fe(II)Cl_2_. The few cells that appeared are likely to be suppressor mutants. On the other hand, the cells grew with the addition of 0.5 mM Fe(III)Cl_3_. Since Fe(III)citrate consists of ferric iron and the iron chelator citrate, we wondered whether the growth phenotype had something to do with the chelation of iron. Intriguingly, the addition of Na_3_citrate (1 g/l) to the plates allows growth of the Δ*furA* mutant also with Fe(II)Cl_2_. Moreover, the addition of sodium citrate to plates without any additional iron also led to weak growth of the *furA* mutant. Taken together, these results indicate that the *furA* is able to grow with ferrous iron sources, provided they are made available by citrate-mediated chelation. In this case, even the low amounts of iron present in the sporulation medium allow at least a faible growth. Thus, the problem with ferrous iron is its low bioavailability.

### FurA affects the activity of the Fur-controlled *dhbA* promoter

The *dhbACEBF-ybdZ* operon encodes the enzymes for the synthesis of the siderophore bacillibactin (36). It is one of the Fur-controlled operons, and it is induced as a part of the second wave of Fur-controlled genes that are expressed upon iron limitation (15). We used the activity of the *dhbA* promoter as an indicator of Fur activity. For this purpose, we fused the *dhbA* promoter region to a promoterless *lacZ* gene encoding β-galactosidase and integrated this *dhbA-lacZ* fusion into the *B. subtilis* genome of the wild type strain and the isogenic Δ*fur* mutant GP879. The resulting strains, GP3331 and GP3356, respectively, carrying the *dhbA-lacZ* fusion were cultivated in CSE-Glc minimal medium supplemented with 0.1 µM and 500 µM of ferrous and ferric iron. As shown in Fig. 2, a strong promoter activity was determined at 0.1 µM iron, irrespective of the redox state of the iron ions. Expression was substantially reduced at the increased iron concentration in the wild type strain. For ferrous iron, we observed an about twofold decrease of expression, whereas a twelvefold decrease was detected in the case of ferric iron. This result corresponds well to the observation that ferrous iron is less bioavailable for the bacteria, so that it causes only a weak repression of *dhbA* expression. In the *fur* mutant GP3356, we observed high β-galactosidase under all tested conditions. The strong repression of *dhbA* expression in response to ferrous iron, and the dependence of repression on a functional Fur protein is in excellent agreement with published data (37). We also tested the expression of the *dhbA-lacZ* fusion in the wild type strain GP3331 in complex medium (LB) in the presence and absence of 500 µM ferrous or ferric iron. In this case, the activity was already rather low in the absence of added iron (45 unit per mg of protein as compared to 700 … 900 units in minimal medium in the presence of 0.1 µM of iron), probably due to the presence of iron in the complex medium which is likely to cause a basal repression. However, the expression was fivefold repressed if one of the iron ions was added (see Fig. 3A for Fe^3+^, data not shown for Fe^2+^). For further experiments, we used LB medium and ferric iron.

**Fig. 2.**
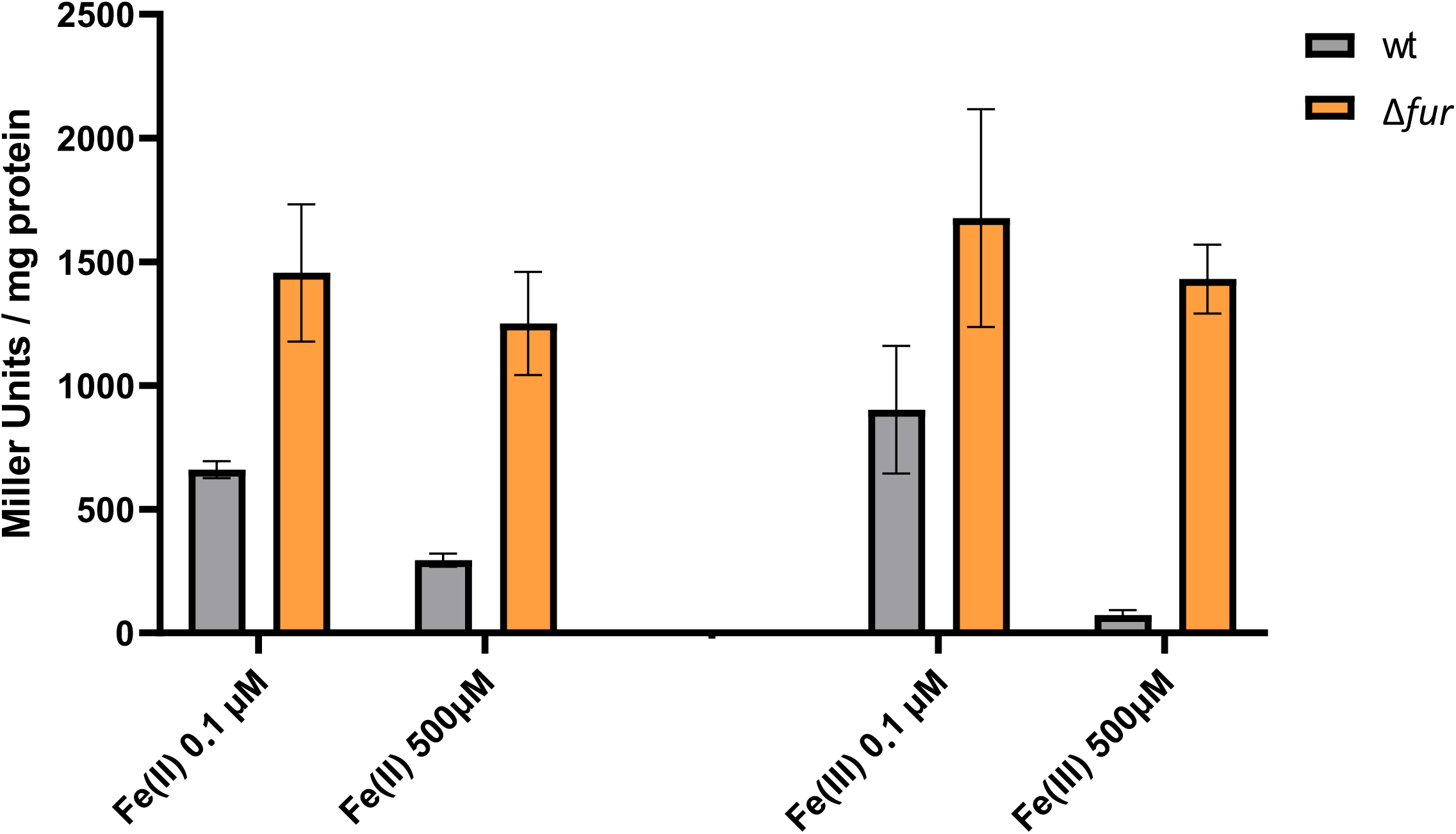
Effect of iron on the activity of the *dhbA* promoter. Cultures of a wild type strain (GP3331) and the isogenic Δ*fur* mutant (GP3356) carrying a *dhbA-lacZ* fusion were grown with the indicated iron supplementation, and promoter activities were determined by quantification of β-galactosidase activities. The values are averages of three independent experiments. Standard deviations are shown.

**Fig. 3.**
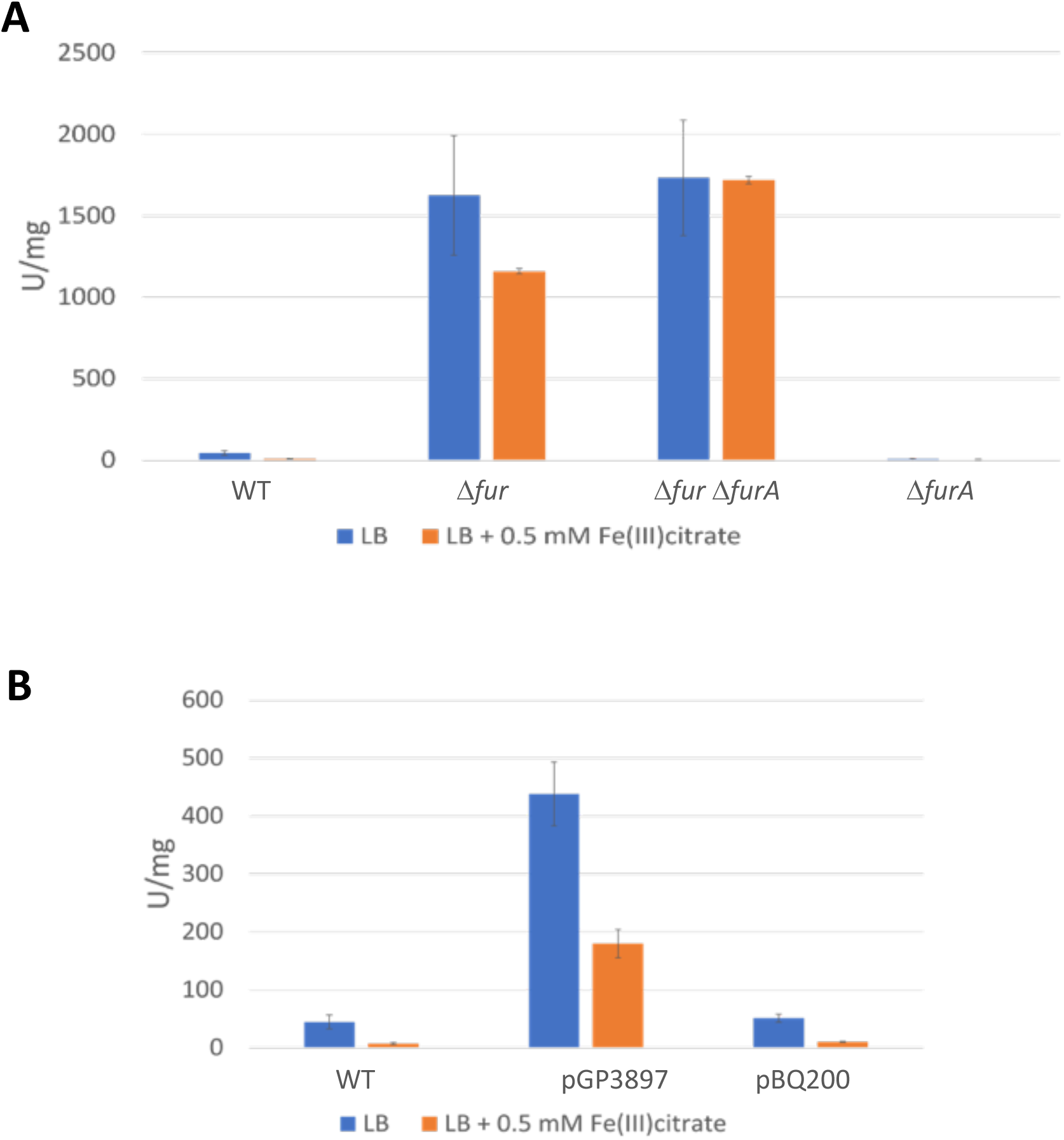
The impact of FurA on *dhbA* promoter activity. A. Effect of *furA* deletion. Strains carrying a *dhbA-lacZ* fusion were cultivated in LB with or without added ferric citrate, and promoter activities were determined by quantification of β-galactosidase activities. The values are averages of three independent experiments. Standard deviations are shown. WT, GP3331; Δ*fur*, GP3356, Δ*fur* Δ*furA*, GP3361; Δ*furA*, GP3366. B. Effect of *furA* overexpression. *B. subtilis* GP3331 without any plasmid (WT) and GP3331 carrying plasmid pGP3897 for *furA* overexpression or the empty vector pBQ200 were cultivated in LB with or without added ferric citrate, and promoter activities were determined by quantification of β-galactosidase activities. The values are averages of three independent experiments. Standard deviations are shown.

To test the role of FurA in the Fur-mediated regulation of the *dhbA* promoter, we constructed additional strains carrying the *dhbA-lacZ* fusion in a *furA* single and *furA fur* double mutant. The resulting strains were GP3366 and GP3361, respectively. As shown in Fig. 3A, expression in the *fur* mutant was strongly enhanced as compared to the wild type background and independent from the iron concentration. The rather weak expression in the wild type strain in the absence of added iron is in good agreement with the conclusion that this medium already contains some iron. The loss of FurA in addition to Fur in GP3361 had no effect on the *dhbA* promoter activity. Finally, we tested *dhbA* promoter activity in the *furA* mutant (GP3366). For this strain, only background activities were observed under both conditions. We conclude that FurA exerts its role via Fur, as already suggested as a result from the suppression analysis. The lack of promoter activity in the *furA* single mutant is in excellent agreement with the above conclusion that the Fur-controlled genes for iron uptake might be completely repressed in the absence of FurA thus resulting in the essentiality of FurA in standard LB medium.

The deletion of *furA* resulted in complete repression of the *dhbA* promoter, likely due to the inability to release Fur from its target DNA in the absence of FurA. Given the observed physical interaction between the two proteins, it is tempting to speculate that the overexpression of FurA might cause the release of Fur from its target sites and thus induction of *dhbA* expression even in the presense of iron. To test this hypothesis, we put the *furA* gene under the control of the strong *degQ*^hy^ promoter in the expression plasmid pBQ200. The resulting plasmid was pGP3897. This plasmid as well as the empty vector pBQ200 were then introduced into strain GP3331 that harbors the *dhbA-lacZ* fusion. Again, the strains were grown in LB medium in the presence or absence of 500 µM ferric citrate. As observed before for the wild type strain, we found a fivefold repression of *dhbA* expression in the presence of the empty vector (see Fig. 3B). In contrast, expression was strongly increased in the presence of pGP3897 when FurA was overproduced. We even observed a substantial expression in this strain if 500 µM ferric citrate were present. This observation indicates that the overexpression of FurA counteracts the repressing effect caused by Fur and strongly suggests that FurA acts as a *bona fide* antagonist of Fur.

### Physical interaction between Fur and FurA

Previous proteome-wide interaction studies with *B. subtillis* detected an interaction between Fur and FurA (33, 34). If FurA acts as an antagonist to Fur, it seems most likely that this activity is achieved by the physical interaction between the two proteins. To confirm this interaction, we decided to verify the interaction *in vivo* in *B. subtilis* and to reconstitute it in a heterologous system.

To confirm the interaction between Fur and FurA, and to distinguish primary from indirect interactions, we studied the binary interaction using the bacterial two-hybrid screen. In this system, interacting proteins reconstitute the *B. pertussis* adenylate cyclase, resulting in cAMP synthesis and subsequent activation of β-galactosidase synthesis. As shown in Fig. 4A, Fur and FurA showed a strong direct interaction. This observation is in excellent agreement with previous results (33, 34). Moreover, FurA exhibited a strong self-interaction which corresponds to the previously reported formation of homodimers for the FurA protein from *Staphylococcus aureus* (38). None of the two proteins showed an interaction with the Zip protein, which was used as the negative control. Thus, the interaction between FurA and Fur is specific. To test, whether the interaction depends on the availability of iron, we also performed the two hybrid screen in the presence of 500 µM of either ferric or ferrous iron. The results were indistiguishable from those obtained in the absence of added iron (data not shown). This may result from internal iron accumulation in *E. coli*.

**Fig. 4.**
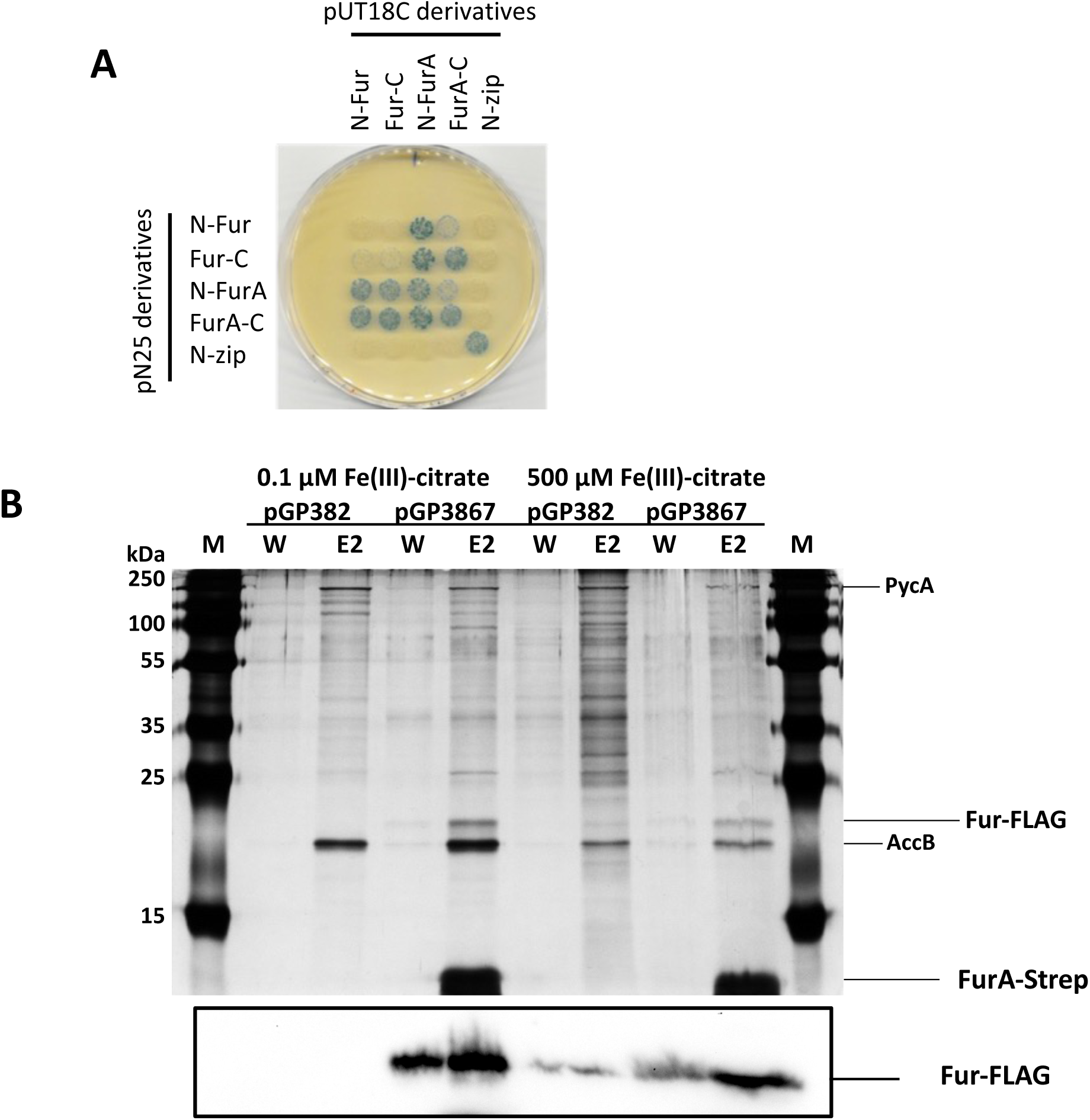
Physical interaction between FurA and Fur. A. Bacterial two-hybrid assay to test the interaction between Fur and FurA. N- and C-terminal fusions of both proteins to the T18 or T25 domain of the adenylate cyclase (CyaA) were created and the proteins were tested for interaction in *E. coli* BTH101. Blue colonies indicate an interaction that results in adenylate cyclase activity and subsequent expression of the reporter β-galactosidase. B. Evaluation of proteins co-purified with Fur. Protein complexes isolated from *B. subtilis* GP3367 (Fur-FLAG) containing either the empty vector pGP382 or pGP3867 (FurA-Strep). The strains were grown in CSE-Glc minimal medium supplemented with ferric citrate as indicated. The wash and the second elution fractions from each purification were loaded onto the SDS-PAA gel and analyzed by silver staining. The positions of the intrinsically biotinylated proteins PycA and AccB as well as of Fur- FLAG and FurA-Strep are shown. The lower panel shows a Western blot using antibodies raised against the FLAG tag to detect the Fur-FLAG protein.

In a second setup, we tested the physical *in vivo* interaction between FurA and Fur by co-precipitation (see Fig. 4B). For this purpose, we constructed a strain that expressed Fur carrying a C-terminal FLAG tag for immunological detection (GP3367). This strain was then transformed either with the empty vector pGP382 (39) or with plasmid pGP3867 for the expression of FurA fused to a C-terminal Strep-tag for affinity chromatography. Both strains were cultivated in CSE-Glc minimal medium with 0.1 or 500 µM ferric citrate. The protein extracts were then passed over a StrepTactin column to bind the FurA-Strep, washed and Strep-tagged proteins with their potential interaction partners were eluted. Two proteins, PycA and AccB, were eluted from the StrepTactin column for both the empty vector control and the strain expressing FurA-Strep. These proteins contain a biotin cofactor that causes binding to the matrix and are good indicators that the experimental setup was suitable. Upon expression of FurA-Strep, we observed copurification of a protein of about 20 kDa which corresponds to FLAG-tagged Fur. The identity of the band was confirmed by a Western blot using antibodied directed against the FLAG tag (see Fig. 5). As observed in the two-hybrid screen, the Fur protein was copurified with FurA both at high and low iron concentrations. Again, the low iron concentration used in this experiment may still be sufficient to allow the interaction between the two proteins.

**Fig. 5.**
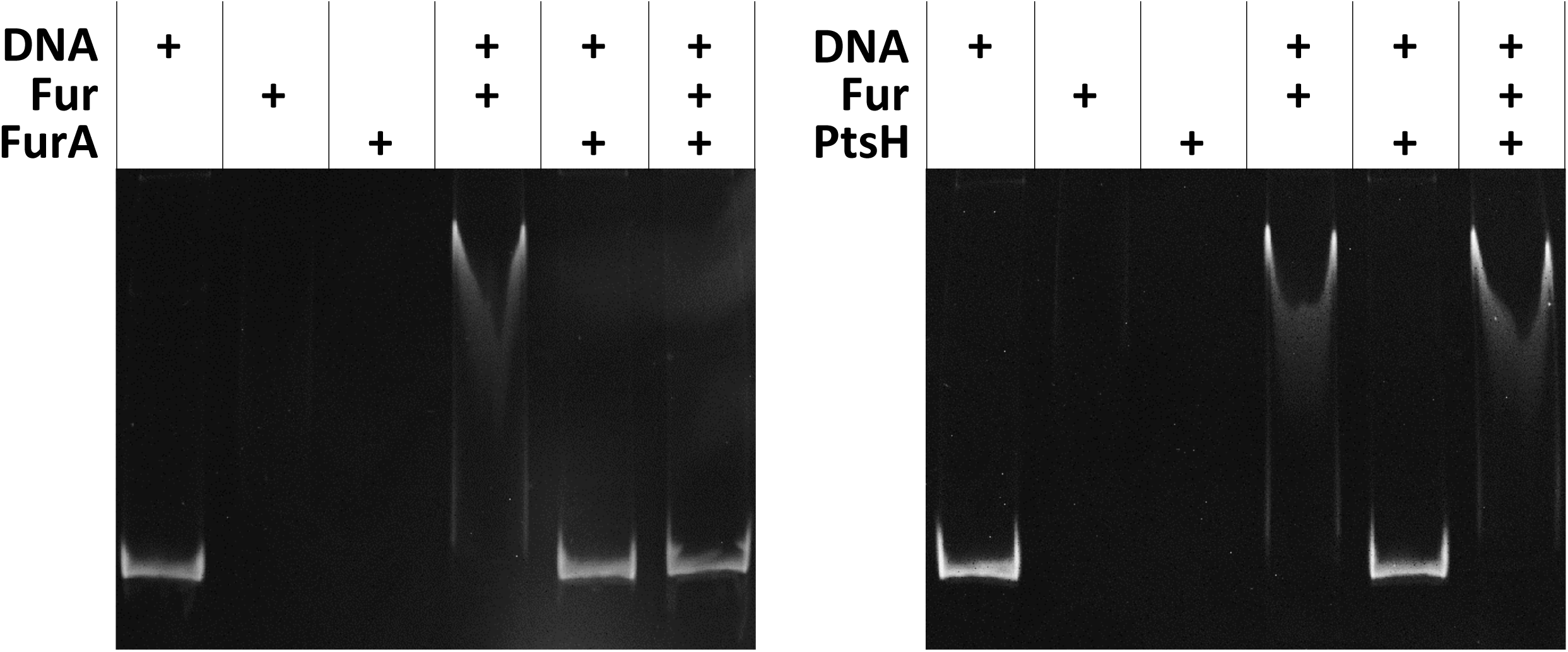
FurA is an antagonist of the DNA-binding activity of Fur. Gel electrophoretic mobility shift assay of Fur binding to *dhbA* promoter fragments. The components added to the assays are shown above the gels.

### FurA prevents the DNA-binding activity of Fur

All our experiments support our initial hypothesis that FurA can bind Fur and interfere with the repression of target genes by Fur. To get direct evidence for this, we performed DNA binding assays with the *dhbA* promoter region and purified Fur protein. As shown in Fig. 5, the DNA fragment was retarded in the presence of Fur, indicating binding of Fur to its target DNA. In contrast, no shift was observed with the purified FurA protein. The addition of both Fur and FurA to the DNA did not result in DNA binding. This observation is in excellent agreement with the idea that FurA might prevent Fur from binding to DNA. To ensure that the effect of FurA addition to the DNA and Fur is specific, we performed a control experiment in which we used the HPr protein of the phosphotransferase system as the seond protein in the assay. As FurA, HPr is a small acidic protein. The results observed with the promoter DNA and Fur were as described above. Similarly, the HPr protein did not interact with the DNA. However, the presence of HPr in addition to the promoter fragment and Fur did not prevent the formation of the Fur-DNA complex indicating that HPr is unable to interfere with the DNA binding activity of Fur (Fig. 5). Taken together, our results demonstrate that FurA specifically interacts with Fur to prevent it from binding to its target DNA sequences and thus to allow the expression of genes that are under negative control by Fur.

### AlphaLink based model for the FurA-Fur complex

The AlphaLink (40) prediction (model confidence: 0.75) of the *B. subtilis* Fur-dimer (Fig. 6A) resembles a V-shaped conformation, similar to other Fur-proteins which interact with DNA (41). If we include the FurA interaction, Fur undergoes a large conformational change (Fig. 6B). The AlphaLink model (model confidence: 0.84) directly leverages the experimental crosslinking MS data (34, 42) which support the interaction. Cross-links were found between Lys-74 in the DNA-binding domain of Fur and the Lys residues 23 and 26 of FurA (34). Upon interaction with Fur, FurA grabs the DNA-binding domains of Fur and the Fur dimerization interface disassembles, resulting in a breaking of the functional dimer and a complete re-orientation of the DNA-binding domain (see http://www.subtiwiki.uni-goettingen.de/v4/predictedComplex?id=168 and http://www.subtiwiki.uni-goettingen.de/v4/predictedComplex?id=169 for an interactive display of the Fur dimer and the Fur-FurA complex as well as Supplementary Movie S1). The rotation and acompanying re-orientation of the DNA-binding domains explains the loss of the DNA-binding activity of Fur upon interaction with FurA.

**Fig. 6.**
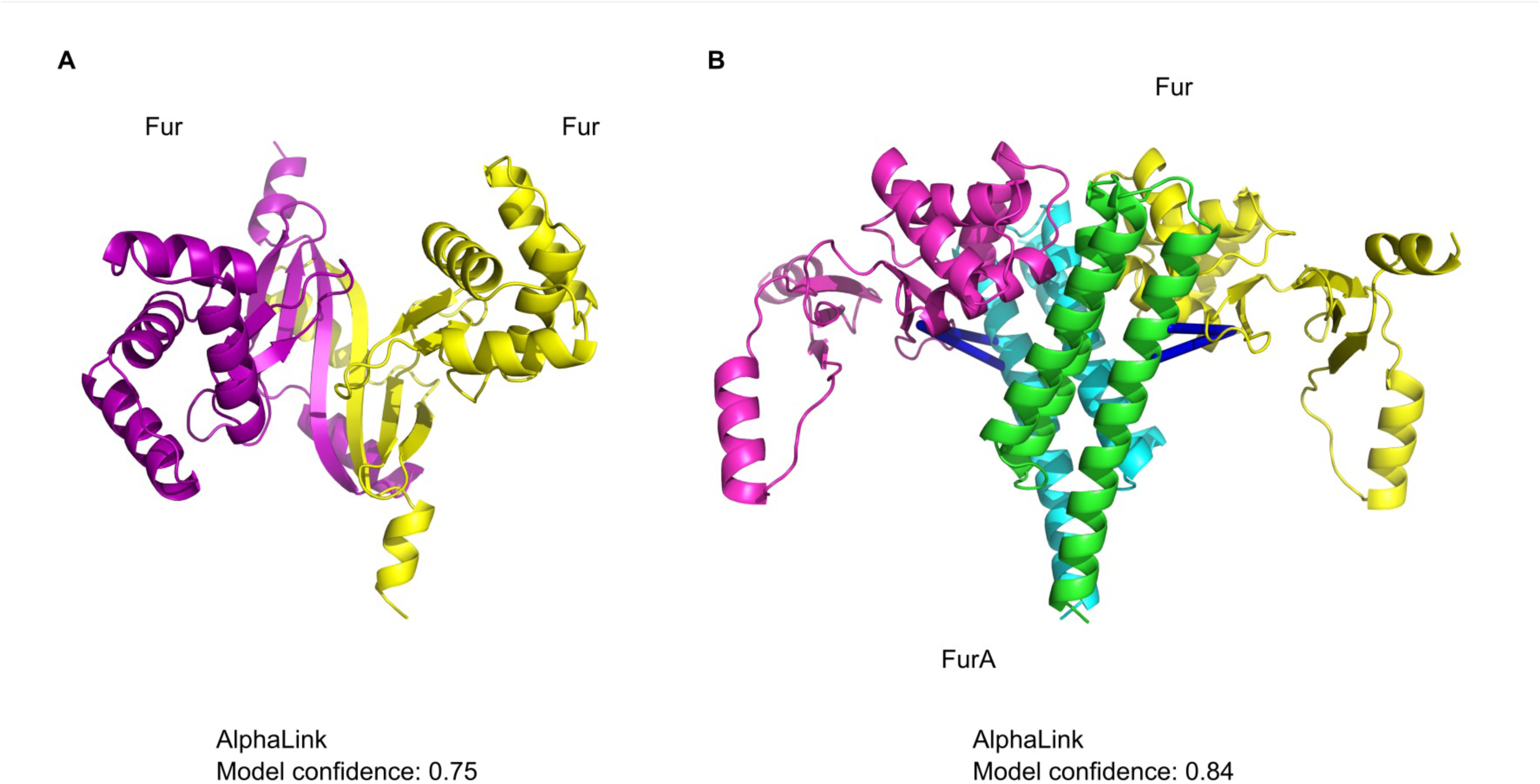
AlphaLink and cross-linking data suggest the structure of the FurA-Fur complex. A: AlphaLink prediction (model confidence: 0.75) of the Fur dimer. **B:** AlphaLink prediction (model confidence: 0.84) of the FurA-Fur complex with crosslinking MS data. The Fur dimer undergoes a large conformational change. All crosslinks (shown as blue lines) are satisfied in the prediction.

## DISCUSSION

The FurA (YlaN) protein belongs to a small group of so far unknown proteins that are strongly expressed under essentially all conditions in *B. subtilis* (29). Of those about 40 proteins, FurA is the only that is essential under standard laboratory growth conditions (30, 35). This made the protein an important target for functional analysis. The data presented in this study identify the so far unknown protein FurA as a *bona fide* antirepressor of the Fur regulator. The interaction between the two proteins interferes with the binding of Fur to its DNA targets and thus results in the expression of the iron uptake systems which are all subject to Fur-mediated transcription repression.

The data presented in the acompanying paper suggest that FurA perceives the primary signal of the system, the intracellular iron concentration in the form of ferrous iron (43). In this form, the protein does not interact with Fur, and Fur can bind to its DNA target sites to repress the expression of genes for iron uptake systems and to activate expression of an iron exporter gene. In contrast, under conditions of iron limitation, apo-FurA can bind to Fur, and thus break open the Fur dimer, resulting in release of Fur from its target sites as shown in this work (see Fig. 7). This interaction allows the expression of Fur-controlled genes for iron uptake systems if iron gets scarce, and is thus a prerequisite for the growth of *B. subtilis* under conditions of iron depletion. This explains why the FurA gene is essential for *B. subtilis* under standard conditions. Since iron limitation is the rule rather than the exception for bacteria that live under aerobic conditions, mechanisms that allow the effective induction of genes for iron acquisition are of key importance.

**Fig. 7.**
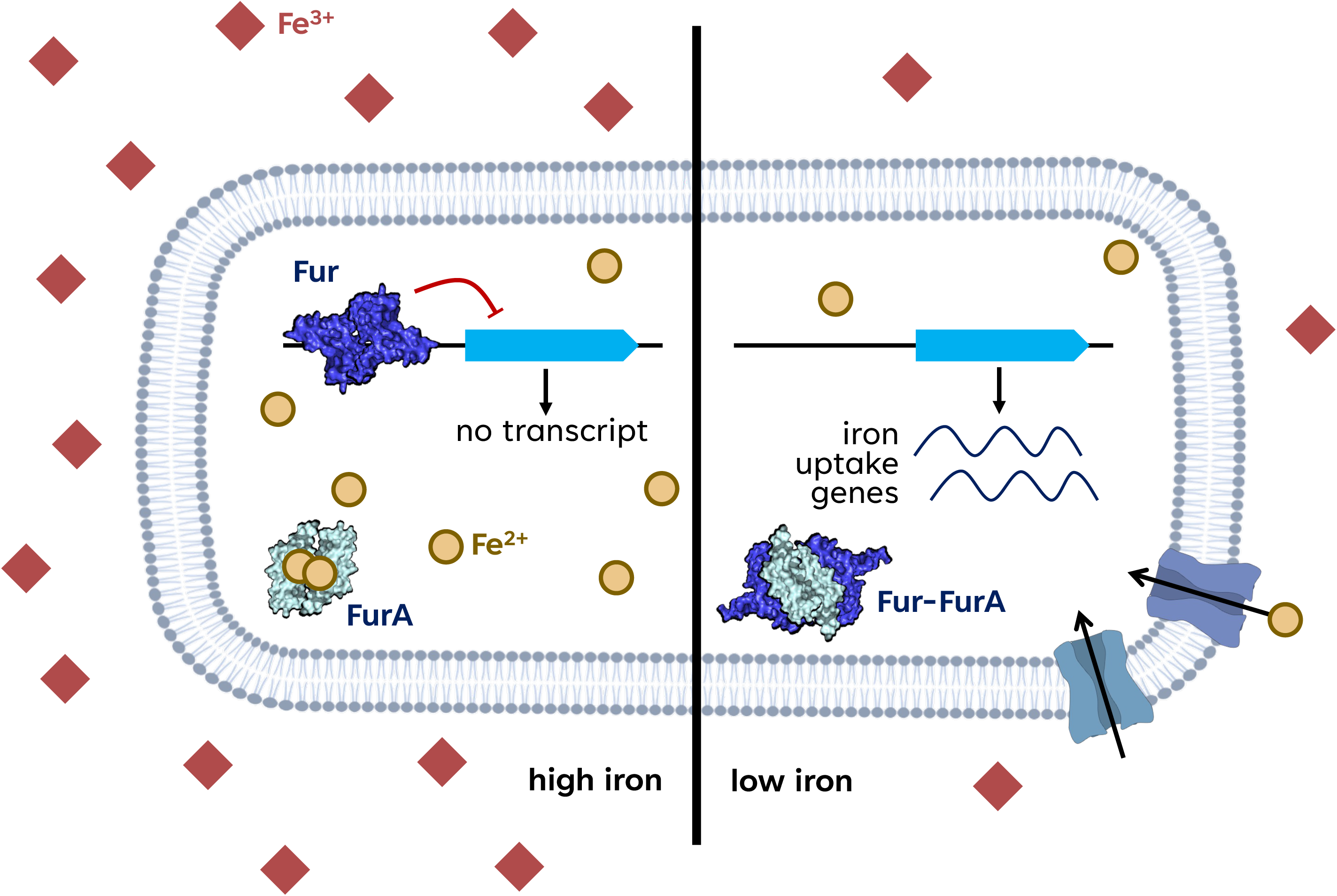
Model for the control of Fut activity by FurA. At high iron concentrations, the Fur dimer binds its target DNA sequences in the promoter regions of genes involved in iron homeostasis. The FurA protein binds ferrous iron and is unable to interact with Fur. If iron gets limiting, apo-FurA forms a complex with Fur, resulting in the release of Fur from its targets, and thus in expression of genes for iron uptake.

Fur-mediated control of iron homeostasis is widespread in both Gram- negative and Gram-positive bacteria. For long time, it was assumed that the Fur protein directly responds to the presence of ferrous iron (44). However, despite intensive research there is no clear support for this idea in the published data body and the direct sensing of iron by Fur has remained controversial (45, 46). In contrast, there is clear evidence that zinc ions act as cofactor for the regulator of zinc homeostasis Zur, another Fur-type regulator (47). Only more recent studies with Gram-negative bacteria and the data presented in this and the acompanying study suggest that Fur may be controlled by regulatory protein-protein interactions in many bacteria (25, 43, for review). As observed in this study, the Fur antagonists YdiV and PtsN of *E. coli* and *S. enterica*, respectively, are required in both bacteria to allow expression of Fur-repressed genes if the cell experience iron limitation (26, 27).

An analysis of the phylogenetic distribution of FurA reveals that the protein is present exclusively in the Bacilli subgroup of the Firmicutes (13). In this class, FurA is present in most species with the exception of the lactic acid bacteria and few other species. Interestingly, most bacteria that possess Fur family transcription factors, encode multiple, typically three of these proteins, Fur, PerR, and Zur. Those bacteria of the Bacilli class that lack FurA, do also lack the Fur protein. Most of them have PerR and Zur, with the notable exception of *Lactobacillus acidophilus* and *Streptococcus pneumoniae* that completely lack Fur type regulators. In the genus *Jeotgalibacillus*, one species, *J. malaysiensis* possesses both Fur and FurA, whereas *J. donkookensis* encodes neither of the two proteins. There are only two bacteria among the Bacilli that seem to possess Fur but not FurA, *Aneurinibacillus soli* CB4 and *Tumebacillus avium* AR23208. This might result from issues with the genome sequences, or these bacteria have evolved specific strategies to control Fur activity.

The two-faced role of iron as an important player in cellular energy metabolism on one hand and its toxicity on the other make the presence of effective systems to control iron homeostasis critical for bacterial life. *B. subtilis* possesses several systems that sense and respond to iron availability. The global system is the Fur/ FurA repressor/ antirepressor couple which controls the expression of iron uptake and export systems, of proteins that counteract oxidative stress, and of a regulatory RNA and its chaperone proteins (15, 16). As a result of Fur/FurA-dependent regulation, the genes for iron uptake are not expressed if the metal is already abundant in the cell. The antirepressor activity of FurA allows expression of iron uptake genes as soon as iron gets limiting. This is a strategic decision of the cell as it determines which protein of the iron homeostasis system will be present or not in the future. However, Fur-dependent control is not sufficient if iron gets limiting and immediate measures must be taken to increase its availability. Citrate chelates iron, suggesting that large citrate pools are rather problematic for the cell if iron gets limiting. Indeed, *B. subtilis* has also found a solution for this problem: aconitase, an enzyme of the citric acid cycle, needs an FeS cofactor. In the absence of iron, apo-aconitase is an RNA-binding protein. In *B. subtilis*, it binds to the *citZ* mRNA that specifies citrate synthase to trigger its degradation, and to prevent the synthesis of even more citrate synthase from pre-synthesized mRNA molecules (48, 49). To be completely on the safe side and to prevent any further citrate synthesis, the cell should also degrade or otherwise inactivate citrase synthase during iron limitation; however, this issue remains to be investigated.

The discovery of the activity of FurA as an antirepressor to Fur in *B. subtilis* is important for our better understanding of the physiology of this important model organism (50). *B. subtilis* is the model organism for a large group of Gram-positive bacteria, that includes many important pathogens such as *S. aureus*, *Listeria monocytogenes*, or *Bacillus anthracis* that also possess the Fur/FurA couple. Iron is the growth-limiting factor for most pathogenic bacteria in the human body. Accordingly, the investigation of the control of iron homeostasis is also very important to better understand the processes of infection and disease and to develop novel treatments. Since FurA is essential under conditions of iron limitation not only in *B. subtilis* but also in *S. aureus* (see acompanying paper, 43), it might be an attractive novel target for drug development.

## MATERIALS AND METHODS

### Strains, media and growth conditions

*E. coli* DH5α and Rosetta DE3 (51) were used for cloning and for the expression of recombinant proteins, respectively. All *B. subtilis* strains used in this study are derivatives of the laboratory strain 168. They are listed in Table 1. *B. subtilis* and *E. coli* were grown in Luria-Bertani (LB) or in sporulation (SP) medium (51,52). For growth assays and the *in vivo* interaction experiments, *B. subtilis* was cultivated in LB, SP, or CSE-Glc minimal medium (52, 53). CSE-Glc is a chemically defined medium that contains sodium succinate (6 g/l), potassium glutamate (8 g/l), and glucose (1 g/l) as the carbon sources (53). Iron sources were added as indicated. The media were supplemented with ampicillin (100 µg/ml), kanamycin (50 µg/ml), chloramphenicol (5 µg/ml), or erythromycin and lincomycin (2 and 25 µg/ml, respectively) if required. LB and SP plates were prepared by the addition of Bacto Agar (Difco) (17 g/l) to the medium.

**Table 1.** *B. subtilis* strains used in this study.

| Strain | Genotype | Source or Reference |
| --- | --- | --- |
| 168 | *trpC2* | Laboratory collection |
| GP879 | *trpC2 Δfur::ermC* | This study |
| GP3321 | *trpC2 Δfur::ermC ΔfurA::cat* | This study |
| GP3324 | *trpC2 ΔfurA::cat* | This study |
| GP3331 | *trpC2 amyE::(P<sub>dhbA</sub>-lacZ aphA3)* | pGP3594 → 168 |
| GP3356 | *trpC2 Δfur::ermC amyE::(P<sub>dhbA</sub>-lacZ aphA3)* | GP879 → GP3331 |
| GP3361 | *trpC2 Δfur::ermC ΔfurA::cat amyE::(P<sub>dhbA</sub>-lacZ aphA3)* | GP3321 → GP3331 |
| GP3366 | *trpC2 ΔfurA::cat amyE::(P<sub>dhbA</sub>-lacZ aphA3)* | GP3324 → GP3331 |
| GP3367 | *trpC2 fur-FLAG spc* | pGP3899 → 168 |
| GP3368 | *trpC2 ΔfurA::cat* point mutations in *spolIM-fur*<br>intergenic region | Suppressor of GP3324 (no<br>iron) |

### DNA manipulation

Transformation of *E. coli* and plasmid DNA extraction were performed using standard procedures (51). All commercially available plasmids, restriction enzymes, T4 DNA ligase and DNA polymerases were used as recommended by the manufacturers. *B. subtilis* was transformed with plasmids, genomic DNA or PCR products according to the two-step protocol (52). Transformants were selected on SP plates containing erythromycin (2 µg/ml) plus lincomycin (25 µg/ml), chloramphenicol (5 µg/ml), kanamycin (10 µg/ml), or spectinomycin (250 µg/ml). DNA fragments were purified using the QIAquick PCR Purification Kit (Qiagen, Hilden, Germany). DNA sequences were determined by the dideoxy chain termination method (51).

### Construction of mutant strains by allelic replacement

Deletion of the *fur* and *furA* genes was achieved by transformation of *B. subtilis* 168 or GP879 with a PCR product constructed using oligonucleotides to amplify DNA fragments flanking the target genes and an appropriate intervening resistance cassette as described previously (54). The integrity of the regions flanking the integrated resistance cassette was verified by sequencing PCR products of about 1,100 bp amplified from chromosomal DNA of the resulting mutant strains.

### Phenotypic analysis

In *B. subtilis*, amylase activity was detected after growth on plates containing nutrient broth (7.5 g/l), 17 g Bacto agar/l (Difco) and 5 g hydrolyzed starch/l (Connaught). Starch degradation was detected by sublimating iodine onto the plates.

Quantitative studies of *lacZ* expression in *B. subtilis* were performed as follows: cells were grown in CSE-Glc or LB medium supplemented with iron sources as indicated as indicated. Cells were harvested at OD_600_ of 0.5 to 0.8. β-Galactosidase specific activities were determined with cell extracts obtained by lysozyme treatment as described previously (52). One unit of β-galactosidase is defined as the amount of enzyme which produces 1 nmol of o-nitrophenol per min at 28° C.

### Genome sequencing

To identify the mutations in the suppressor mutant strains GP3368 (see Table 1), the genomic DNA was subjected to whole-genome sequencing. Concentration and purity of the isolated DNA was first checked with a Nanodrop ND-1000 (PeqLab Erlangen, Germany), and the precise concentration was determined using the Qubit® dsDNA HS Assay Kit as recommended by the manufacturer (Life Technologies GmbH, Darmstadt, Germany). Illumina shotgun libraries were prepared using the Nextera XT DNA Sample Preparation Kit and subsequently sequenced on a MiSeq system with the reagent kit v3 with 600 cycles (Illumina, San Diego, CA, USA) as recommended by the manufacturer. The reads were mapped on the reference genome of *B. subtilis* 168 (GenBank accession number: NC_000964) (55). Mapping of the reads was performed using the Geneious software package (Biomatters Ltd., New Zealand) (56). Frequently occurring hitchhiker mutations (57) and silent mutations were omitted from the screen.

The resulting genome sequence was compared to that of our in-house wild type strain. Single nucleotide polymorphisms were considered as significant when the total coverage depth exceeded 25 reads with a variant frequency of ≥90%. All identified mutations were verified by PCR amplification and Sanger sequencing.

### Plasmid constructions

To express the Fur and FurA proteins carrying a N-terminal His-tag in *E. coli*, the *fur* and *furA* genes were amplified using chromosomal DNA of *B. subtilis* 168 as the template and appropriate oligonucleotides that attached specific restriction sites to the fragment. Those were: BamHI and XhoI for cloning *fur* in pET-SUMO (Invitrogen, Germany), and BamHI and SalI for cloning *furA* in pWH844 (58). The resulting plasmids were pGP3589 and pGP2583 for Fur and FurA, respectively.

For overexpression of *furA* in *B. subtilis*, we constructed plasmid pGP3897. For this purpose, the *furA* gene was amplified and cloned between the BamHI and SalI site of the expression vector pBQ200 (59). For the expression of FurA carrying a C-terminal Strep-tag in *B. subtilis*, we used plasmid pGP3867.

This plasmid was obtained by cloning the *furA* gene between the BamHI and SalI sites of the expression vector pGP382 (39). To add a FLAG tag epitope to the Fur protein, we constructed plasmid pGP3899 by cloning the fur gene between the BamHI and HindIII sites of pGP1331 (60).

Plasmid pAC7 (61) was used to a construct translational fusion of the *dhbA* promoter region to the promoterless *lacZ* gene. For this purpose, the promoter region was amplified using oligonucleotides that attached EcoRI and BamHI restriction to the ends of the products. The fragments were cloned between the EcoRI and BamHI sites of pAC7. The resulting plasmid was pGP3594.

### Protein expression and purification

*E. coli* Rosetta(DE3) was transformed with the plasmid pGP371 (62), pGP2583, and pGP3589 encoding His-tagged versions of PtsH, FurA, and Fur, respectively. For overexpression, cells were grown in 2x LB and expression was induced by the addition of isopropyl 1- thio-β-D-galactopyranoside (final concentration, 1 mM) to exponentially growing cultures (OD_600_ of 0.8). The His-tagged proteins were purified in 1x ZAP buffer (50 mM Tris-HCl, 200 mM NaCl, pH 7.5). Cells were lysed by four passes (18,000 p.s.i.) through an HTU DIGI-F press (G. Heinemann, Germany). After lysis, the crude extract was centrifuged at 46,400 x g for 60 min and then passed over a Ni^2+^nitrilotriacetic acid column (IBA, Göttingen, Germany). The proteins were eluted with an imidazole gradient. After elution, the fractions were tested for the desired protein using SDS-PAGE. The purified proteins were concentrated in a Vivaspin turbo 15 (Sartorius) centrifugal filter device (cut-off 5 or 50 kDa). The protein samples were stored at −80°C until further use. The protein concentration was determined according to the method of Bradford (63) using the Bio-Rad dye binding assay and bovine serum albumin as the standard.

### Electromobility shift assay (EMSA) with DNA

To analyze the binding of Fur to the *dhbA* promoter region, we performed EMSA assays with a 284 bp *dhbA* promoter fragment that carries the Fur binding site and purified Fur, FurA, and PtsH proteins. 200 ng of DNA and 80 pmoI of the proteins were used. The samples were first prepared without the proteins only with DNA, buffer and water and heated for 2 minutes at 95°C. Then the proteins were added in different combinations and the samples were incubated for 30 minutes at 37°C. Meanwhile, the EMSA gels were applied to a pre run at 90 V for 30 minutes immersed in TBE buffer (51). Afterwards, 2 µl of the loading dye were added and the samples were loaded into the gel pockets. The gel was run for 3 hours at 110 V. Then, the gels were immersed in TBE containing HDGreen® fluoreszence dye (Intas, Germany). After 2 minutes the gels were photographed under UV light.

### *In vivo* detection of protein-protein interactions

To verify the interaction between Fur and FurA *in vivo*, cultures of *B. subtilis* GP3367 (Fur-FLAG) containing pGP3867 (FurA-Strep), or the empty vector control (pGP382), were cultivated in 500 ml CSE-Glc medium containing the indicated iron source until exponential growth phase was reached (OD_600_ ∼ 0.4-0.6). The cells were harvested immediately and stored at -20°C. The Strep-tagged protein and its potential interaction partners were then purified from crude extracts using a StrepTactin column (IBA, Göttingen, Germany) and D-desthiobiotin as the eluent. The eluted proteins were separated on an SDS gel and potential interacting partners were analyzed by staining with Colloidal Coomassie and Western blot analysis using antibodies raised against the FLAG-tag.

### Bacterial two-hybrid assay

Primary protein-protein interactions were identified by bacterial two-hybrid (BACTH) analysis (64). The BACTH system is based on the interaction-mediated reconstruction of *Bordetella pertussis* adenylate cyclase (CyaA) activity in *E. coli* BTH101. Functional complementation between two fragments (T18 and T25) of CyaA as a consequence of the interaction between bait and prey molecules results in the synthesis of cAMP, which is monitored by measuring the β-galactosidase activity of the cAMP-CAP-dependent promoter of the *E. coli lac* operon. Plasmids pUT18C and p25N allow the expression of proteins fused to the T18 and T25 fragments of CyaA, respectively. For these experiments, we used the plasmids pGP3868-pGP3875, which encode N-and C-terminal fusions of T18 or T25 to *fur* and *furA.* The plasmids were obtained by cloning the *fur* and *furA* between the KpnI and BamHI sites of pUT18C and p25N (64). The resulting plasmids were then used for co-transformation of *E. coli* BTH101 and the protein-protein interactions were then analyzed by plating the cells on LB plates containing 100 µg/ml ampicillin, 50 µg/ml kanamycin, 40 µg/ml X-Gal (5-bromo-4-chloro-3-indolyl-ß-D-galactopyranoside), and 0.5 mM IPTG (isopropyl-ß-D-thiogalactopyranoside). The plates were incubated for a maximum of 36 h at 28°C.

### Modelling of the structure of the FurA-Fur complex

We predicted the FurA-Fur complex structure with AlphaLink2 v2 (40) using crosslinking MS data (34). For each prediction, we ran 3 recycling iterations. The final model is the best prediction out of 5 samples picked by highest model confidence.

## ACKNOWLEDGEMENTS

We wish to thank Christina Herzberg for support in the lab and Hinnerk Eilers for his help with strain construction. Christoph Elfmann is acknowledged for the help with preparing Movie S1.

## Author contributions

Design of the study: R.B. and J.S. Experimental work: L.D., R.B., B.H. Data analysis: L.D., R.B., K.S., J.R. and J.S. Visualization: L.D., R.B., B.H., K.S., and R.W. Wrote the paper: L.D., R.B., K.S., R.W., J.R. and J.S.

## Notes

### Competing Interest Statement

The authors have declared no competing interest.

## REFERENCES

1. Andrews SC, Robinson AK, Rodríguez-Quiñones F. 2003 Bacterial iron homeostasis. FEMS Microbiol Rev. 27:215–237.

2. Imlay JA. 2003. Pathways of oxidative damage. Annu Rev Microbiol. 57:395–418.

3. Winterbourn CC. 1995. Toxicity of iron and hydrogen peroxide: the Fenton reaction. Toxicol Lett. 82-83:969–974.

4. Kehrer JP. 2000. The Haber-Weiss reaction and mechanisms of toxicity. Toxicology. 149:43–50.

5. Park S, You X, Imlay JA. 2005. Substantial DNA damage from submicromolar intracellular hydrogen peroxide detected in Hpx- mutants of *Escherichia coli*. Proc Natl Acad Sci USA 102:9317–9322.

6. Camprubi E, Jordan SF, Vasiliadou R, Lane N. 2017. Iron catalysis at the origin of life. IUBMB Life. 69:373–381.

7. Williams RJP. 2012. Iron in evolution. FEBS Lett. 586:479–484.

8. Köster W. 2001. ABC transporter-mediated uptake of iron, siderophores, heme and vitamin B12. Res Microbiol. 152:291–301.

9. Ollinger J, Song KB, Antelmann H, Hecker M, Helmann JD. 2006. Role of the Fur regulon in iron transport in *Bacillus subtilis*. J Bacteriol. 188:3664–3673.

10. Stojiljkovic I, Cobeljic M, Hantke K. 1993. *Escherichia coli* K-12 ferrous iron uptake mutants are impaired in their ability to colonize the mouse intestine. FEMS Microbiol Lett 108111–115.

11. Tsolis RM, Bäumler AJ, Heffron F, Stojiljkovic I. 1996. Contribution of TonB- and Feo-mediated iron uptake to growth of *Salmonella typhimurium* in the mouse. Infect Immun. 64:4549–4556.

12. Miethke M, Monteferrante CG, Marahiel MA, van Dijl JM. 2013. The *Bacillus subtilis* EfeUOB transporter is essential for high-affinity acquisition of ferrous and ferric iron. Biochim Biophys Acta. 1833:2267–2278.

13. Galperin MY, Wolf YI, Makarova KS, Vera Alvarez R, Landsman D, Koonin EV. 2021. COG database update: focus on microbial diversity, model organisms, and widespread pathogens. Nucleic Acids Res. 49:D274–D281.

14. Baichoo N, Wang T, Ye R, Helmann JD. 2002. Global analysis of the *Bacillus subtilis fur* regulon and the iron starvation stimulon. Mol Microbiol. 45:1613–1629.

15. Pi H, Helmann JD. 2017. Sequential induction of Fur-regulated genes in response to iron limitation in *Bacillus subtilis*. Proc Natl Acad Sci USA 114:12785–12790.

16. Pedreira T, Elfmann C, Stülke J. 2022. The current state of *Subti*Wiki, the database for the model organism *Bacillus subtilis*. Nucleic Acids Res. 50:D875–D882.

17. Guan G, Pinochet-Barros A, Gaballa A, Patel SJ, Argüello JM., Helmann JD. 2015. PfeT, a P1B4-type ATPase, effluxes ferrous iron and protects *Bacillus subtilis* against iron intoxication. Mol Microbiol. 98:787–803.

18. Pinochet-Barros A, Helmann JD. 2020. *Bacillus subtilis* Fur is a transcriptional activator for the PerR-repressed *pfeT* gene, encoding an iron efflux pump. J Bacteriol. 202:e00697–19.

19. Coy M, Neilands JB. 1991. Structural dynamics and functional domains of the Fur protein. Biochemistry 30:8201–8210.

20. Stojiljkovic I, Hantke K. 1995. Functional domains of the *Escherichia coli* ferric uptake regulator protein (Fur). MGG Mol Gen Genet. 247:199–205.

21. Fontenot CR, Tasnim H, Valdes KA, Popescu CV, Ding H. 2020. Ferric uptake regulator (Fur) reversibly binds a [2Fe-2S] cluster to sense intracellular iron homeostasis in *Escherichia coli*. J Biol Chem. 295:15454–15463.

22. Pinochet-Barros A, Helmann JD. 2017. Redox sensing by Fe^2+^ in bacterial Fur family metalloregulators. Antioxid Redox Signal. 29:1858–1871.

23. Helmann JD. 2014. Specificity of metal sensing: iron and manganese homeostasis in *Bacillus subtilis*. J Biol Chem. 289:28112–28120.

24. Gaballa A, Antelmann H, Aguilar C, Khakh SK, Song KB, Smaldone GT, Helmann JD. 2008. The *Bacillus subtilis* iron-sparing response is mediated by a Fur-regulated small RNA and three small, basic proteins. Proc Natl Acad Sci USA 105:11927–11932.

25. Steingard CH, Helmann JD. 2023. Meddling with metal sensors: Fur-family proteins as signaling hubs. J Bacteriol. 205:e00022–23.

26. Choi J, Ryu S. 2019. Regulation of iron uptake by fine-tuning the iron responsiveness of the iron sensor Fur. Appl Environ Microbiol. 85:e03026–18.

27. Zhang F, Li B, Dong H, Chen M, Yao S, Li J, Zhang H, Liu X, Wang H, Song N, Zhang K, Du N, Xu S, Gu L. 2020. YdiV regulates *Escherichia coli* ferric uptake by manipulating the DNA-binding ability of Fur in a SlyD-dependent manner. Nucleic Acids Res. 48:9571–9588.

28. Nicolas P, Mäder U, Dervyn E, Rochat T, Leduc A, Pigeonneau N, Bidnenko E, Marchadier E, Hoebeke M, Aymerich S, Becher D, Bisicchia P, Botella E, Delumeau O, Doherty G, Denham EL, Fogg MJ, Fromion V, Goelzer A, Hansen A, Härtig E, Harwood CR, Homuth G, Jarmer H, Jules M, Klipp E, Chat LL, Lecointe F, Lewis P, Liebermeister W, March A, Mars RAT, Nannapaneni P, Noone D, Pohl S, Rinn B, Rügheimer F, Sappa PK, Samson F, Schaffer M, Schwikowski B, Steil L, Stülke J, Wiegert T, Devine KM, Wilkinson AJ, Dijl JM van, Hecker M, Völker U, Bessières P, Noirot P. 2012. Condition- dependent transcriptome reveals high-level regulatory architecture in *Bacillus subtilis*. Science 335:1103–1106.

29. Wicke D, Meißner J, Warneke R, Elfmann C, Stülke K. 2023. Understudied proteins and understudied functions in the model bacterium *Bacillus subtilis* – A major challenge in current research. Mol Microbiol. 120:8–19.

30. Hunt A, Rawlins JP, Thomaides HB., Errington J. 2006. Functional analysis of 11 putative essential genes in *Bacillus subtilis*. Microbiology 152:2895–2907.

31. Commichau FM, Pietack N, Stülke J. 2013. Essential genes in *Bacillus subtilis*: a re-evaluation after ten years. Mol Biosyst. 9:1068–1075.

32. Peters JM, Colavin A, Shi H, Czarny TL, Larson MH, Wong S, Hawkins JS, Lu CHS, Koo BM, Marta E, Shiver AL, Whitehead EH, Weissman JS, Brown ED, Qi LS, Huang KC, Gross CA. 2016. A comprehensive, CRISPR-based functional analysis of essential genes in bacteria. Cell 165:1493– 1506.

33. de Jong L, Roseboom W, Kramer G. 2021. A composite filter for low FDR of protein-protein interactions detected by in vivo cross-linking. J Proteomics 230:103987.

34. O’Reilly FJ, Graziadei A, Forbrig C, Bremenkamp R, Charles K, Lenz S, Elfmann C, Fischer L, Stülke J, Rappsilber J. 2023. Protein complexes in cells by AI-assisted structural proteomics. Mol Syst Biol. 19:e11544.

35. Kobayashi K, Ehrlich SD, Albertini A, Amati G, Andersen KK, Arnaud M, Asai K, Ashikaga S, Aymerich S, Bessieres P, Boland F, Brignell SC, Bron S, Bunai K, Chapuis J, Christiansen LC, Danchin A, Débarbouille M, Dervyn E, Deuerling E, Devine K, Devine SK, Dreesen O, Errington J, Fillinger S, Foster SJ, Fujita Y, Galizzi A, Gardan R, Eschevins C, Fukushima T, Haga K, Harwood CR, Hecker M, Hosoya D, Hullo MF, Kakeshita H, Karamata D, Kasahara Y, Kawamura F, Koga K, Koski P, Kuwana R, Imamura D, Ishimaru M, Ishikawa S, Ishio I, Le Coq D, Masson A, Mauël C, Meima R, Mellado RP, Moir A, Moriya S, Nagakawa E, Nanamiya H, Nakai S, Nygaard P, Ogura M, Ohanan T, O’Reilly M, O’Rourke M, Pragai Z, Pooley HM, Rapoport G, Rawlins JP, Rivas LA, Rivolta C, Sadaie A, Sadaie Y, Sarvas M, Sato T, Saxild HH, Scanlan E, Schumann W, Seegers JF, Sekiguchi J, Sekowska A, Séror SJ, Simon M, Stragier P, Studer R, Takamatsu H, Tanaka T, Takeuchi M, Thomaides HB, Vagner V, van Dijl JM, Watabe K, Wipat A, Yamamoto H, Yamamoto M, Yamamoto Y, Yamane K, Yata K, Yoshida K, Yoshikawa H, Zuber U, Ogasawara N. 2003. Essential Bacillus subtilis genes. Proc Natl Acad Sci USA 100:4678–4683.

36. May JJ, Wendrich TM, Marahiel MA. 2001. The *dhb* operon of *Bacillus subtilis* encodes the biosynthetic template for the catecholic siderophore 2,3-dihydroxybenzoate-glycine-threonine trimeric ester bacillibactin. J Biol Chem. 276:7209–7217.

37. Bsat N, Helmann JD. 1999. Interaction of *Bacillus subtilis* Fur (ferric uptake repressor) with the *dhb* operator *in vitro* and *in vivo*. J Bacteriol. 181:4299–4307.

38. Xu L, Sedelnikova SE, Baker PJ, Hunt A, Errington J, Rice DW. 2007. Crystal structure of *S. aureus* YlaN, an essential leucine rich protein involved in the control of cell shape. Proteins. 68:438–445.

39. Herzberg C, Weidinger LAF, Dörrbecker B, Hübner S, Stülke J, Commichau FM. 2007. SPINE: A method for the rapid detection and analysis of protein-protein interactions *in vivo*. Proteomics. 7:4032–4035.

40. Stahl K, Brock O, Rappsilber J. 2023. Modelling protein complexes with crosslinking mass spectrometry and deep learning. bioRxiv, 10.1101/2023.06.07.544059.

41. Butcher J, Sarvan S, Brunzelle JS, Couture JF, Stintzi A. 2012. Structure and regulon of *Campylobacter jejuni* ferric uptake regulator Fur define apo-Fur regulation. Proc Natl Acad Sci USA 109:10047–10052.

42. Elfmann C, Stülke J. 2023. PAE viewer: a webserver for the interactive vizualization of the predicted aligned error for multimer structure predictions and crosslinks. Nucleic Acids Res. 51:W404–W410.

43. Boyd JM, Esquilin-Lebrón K, Campbell CJ, Kaler KR, Norambuena J, Foley ME, Stephens TG, Rios G, Mereddy G, Zheng V, Bovermann H, Kim J, Kulczyk AW, Yan JH, Greco TM, Cristea IM, Carabetta VJ, Beavers WN, Bhattacharya D, Skaar EP, Parker D, Carroll RK, Stemmler TL. 2023. YlaN is an ionic iron binding protein that functions to relieve Fur-mediated repression of gene expression in *Staphylococcus aureus*. Subm.

44. Troxell B, Hassan HM. 2013: Transcriptional regulation by ferric uptake regulator (Fur) in pathogenic bacteria. Front Cell Infect Microbiol. 3:59.

45. Ma Z, Faulkner MJ, Helmann JD. 2012. Origins of specificity and cross-talk in metal ion sensing by *Bacillus subtilis* Fur. Mol Microbiol. 86:1144–1155.

46. Chandrangsu P, Rensing C, Helmann JD. 2017. Metal homeostasis and resistance in bacteria. Nat Rev Microbiol. 15:338–350.

47. Ma Z, Gabriel SE, Helmann JD. 2011. Sequential binding and sensing of Zn(II) by *Bacillus subtilis* Zur. Nucleic Acids Res. 39:9130–9138.

48. Alén C, Sonenshein AL. 1999. *Bacillus subtilis* aconitase is an RNA-binding protein. Proc Natl Acad Sci USA 96:10412–10417.

49. Pechter KB, Meyer FM, Serio AW, Stülke J, Sonenshein AL. 2013. Two roles for aconitase in the regulation of ticarboxylic acid branch gene expression in *Bacillus subtilis*. J Bacteriol. 195:1525–1537.

50. Stülke J, Grüppen A, Bramkamp M, Pelzer S. 2023. *Bacillus subtilis*, a swiss army knife in science and biotechnology. J Bacteriol. 205:e00102–23.

51. Sambrook J, Fritsch EF, Maniatis T. 1989. Molecular cloning: a laboratory manual, 2nd ed. Cold Spring Harbor Laboratory, Cold Spring Harbor, N.Y.

52. Kunst F, Rapoport G. 1995.Salt stress is an environmental signal affecting degradative enzyme synthesis in *Bacillus subtilis*. J Bacteriol 177:2403–2407.

53. Schmalisch MH, Bachem S, Stülke J. 2003. Control of the *Bacillus subtilis* antiterminator protein GlcT by phosphorylation. Elucidation of the phosphorylation leading to the inactivation of GlcT. J Biol Chem. 278:51108–51115.

54. Diethmaier C, Pietack N, Gunka K, Wrede C, Lehnik-Habrink M, Herzberg C, Hübner S, Stülke J. 2011. A novel factor controlling bistability in *Bacillus subtilis*: the YmdB protein affects flagellin expression and biofilm formation. J Bacteriol 193:5997–6007.

55. Barbe V, Cruveiller S, Kunst F, Lenoble P, Meurice G, Sekowska A, Vallenet D, Wang T, Moszer I, Médigue C, Danchin A. 2009. From a consortium sequence to a unified sequence: the *Bacillus subtilis* 168 reference genome a decade later. Microbiology 155:1758–1775.

56. Kearse M, Moir R, Wilson A, Stones-Havas S, Cheung M, Sturrock S, Buxton S, Cooper A, Markowitz S, Duran C, Thierer T, Ashton B, Meintjes P, Drummond A. 2012. Geneious basic: an integrated and extendable desktop software platform for the organization and analysis of sequence data. Bioinformatics 28:1647–1649.

57. Reuß DR, Faßhauer P, Mroch PJ, Ul-Haq I, Koo BM, Pöhlein A, Gross CA, Daniel R, Brantl S, Stülke J. 2019. Topoisomerase IV can functionally replace all type 1A topoisomerases in *Bacillus subtilis*. Nucleic Acids Res. 47:5231–5242.

58. Schirmer F, Ehrt S, Hillen W. 1997. Expression, inducer spectrum, domain structure, and function of MopR, the regulator of phenol degradation in *Acinetobacter calcoaceticus* NCIB8250. J Bacteriol 179:1329–1336.

59. Martin-Verstraete I, Débarbouillé M, Rapoport G. 1994. Interactions of wild-type and truncated LevR of Bacillus subtilis with the upstream activating sequence of the levanase operon. J Mol Biol. 241:178–192.

60. Lehnik-Habrink M, Pförtner H, Rempeters L, Pietack N, Herzberg C, Stülke J. 2010. The RNA degradosome in *Bacillus subtilis*: Identification of Csha as the major RNA helicase in the multiprotein complex. Mol Microbiol. 77:958–971.

61. Weinrauch Y, Msadek T, Kunst F, Dubnau D. 1991. Sequence and properties of *comQ*, a new competence regulatory gene of *Bacillus subtilis*. J Bacteriol. 173:5685–5693.

62. Pietack N, Becher D, Schmidl SR, Saier MH, Hecker M, Commichau FM, Stülke J. In vitro phosphorylation of key metabolic enzymes from Bacillus subtilis: PrkC phosphorylates enzymes from different branches of basic metabolism. J Mol Microbiol Biotechnol 18:129–140.

63. Bradford MM. 1976. A rapid and sensitive method for the quantification of microgram quantities of protein utilizing the principle of protein-dye binding. Anal Biochem 72:248–254.

64. Karimova G, Pidoux J, Ullmann A, Ladant D. 1998. A bacterial two-hybrid system based on a reconstituted signal transduction pathway. Proc Natl Acad Sci USA 95:5752–5756.

