## Supplementary material for "Control of iron homeostasis by a regulatory protein-protein interaction in *Bacillus subtilis*: The FurA (YlaN) acts as an antirepressor to the ferric uptake regulator Fur": Fig. S1

```
                                                                                *
TCATCTCTCT TTGAAGCATA -CTTATCTCC TGTTTTAATG GAAAAGCTTA CATCTAGACT TTTTTAAAAT
TCATCTCTCT TTGAAGCATA GCGTATTT-- TGTTATAATG AGTCATGGAA AACCTAGACT TTGGTAAAAT

                                                      -35
CATTATTAAA AGAAAACCAT TTTATTTATC AGTTTATAAT AATTATAGTT GGAACTCTGC GCGTATTTTG
GCCTTTGTGC TGCTGGTCAT TCTGTTTTTA GCGCTGATTT CATCTCTCTT TGAA-GCATA GCGTATTTTG

    -10
TTATAATGAG TCATGGAATG CGGCGTAGGA GGGAAAGACA TG
TTATAATGAG TCATGGAATG CGGCGTAGGA GGGAAAGACA TG
```

**Fig. S1. Mutations in the *spoIIM-fur* intergenic region of the suppressor mutant GP3368.** The wild type sequence is shown in bold, below the sequence found in GP3368.The stop of the *spoIIM* open reading frame is shown by an asterisk. Mutations are highlighted in red. The -35 and -10 regions of teh fur promoter are underlined and labelled. The start codon of the *fur* open reading frame is hihglighed in green.
